# Alignathon: A competitive assessment of whole genome alignment methods

**DOI:** 10.1101/003285

**Authors:** Dent Earl, Ngan Nguyen, Glenn Hickey, Robert S. Harris, Stephen Fitzgerald, Kathryn Beal, Igor Seledtsov, Vladimir Molodtsov, Brian J. Raney, Hiram Clawson, Jaebum Kim, Carsten Kemena, Jia-Ming Chang, Ionas Erb, Alexander Poliakov, Minmei Hou, Javier Herrero, Victor Solovyev, Aaron E. Darling, Jian Ma, Cedric Notredame, Michael Brudno, Inna Dubchak, David Haussler, Benedict Paten

## Abstract

**Background:** Multiple sequence alignments (MSAs) are a prerequisite for a wide variety of evolutionary analyses. Published assessments and benchmark datasets for protein and, to a lesser extent, global nucleotide MSAs are available, but less effort has been made to establish benchmarks in the more general problem of whole genome alignment (WGA).

**Results:** Using the same model as the successful Assemblathon competitions, we organized a competitive evaluation in which teams submitted their alignments, and assessments were performed collectively after all the submissions were received. Three datasets were used: two of simulated primate and mammalian phylogenies, and one of 20 real fly genomes. In total 35 submissions were assessed, submitted by ten teams using 12 different alignment pipelines.

**Conclusions:** We found agreement between independent simulation-based and statistical assessments, indicating that there are substantial accuracy differences between contemporary alignment tools. We saw considerable difference in the alignment quality of differently annotated regions, and found few tools aligned the duplications analysed. We found many tools worked well at shorter evolutionary distances, but fewer performed competitively at longer distances. We provide all datasets, submissions and assessment programs for further study, and provide, as a resource for future benchmarking, a convenient repository of code and data for reproducing the simulation assessments.

## Introduction

Given a set of sequences, a multiple sequence alignment (MSA) is a partitioning of the residues in the sequences, be they amino-acids or nucleotides, into sets of related residues. Here we are interested in the relationship of evolutionary homology. In some other contexts residues may be aligned with a different aim, as in structural alignments, where residues are aligned if located at the same point in a shared crystal structure superposed on the sequences, even though such structural positions may not necessarily be homologous due to insertions or deletions (Kolodny et al. 2005).

MSA is a fundamental problem in biological sequence analysis because it is a prerequisite for most phylogenetic and evolutionary analyses (Felsenstein 2003, Wallace et al. 2005, Edgar and Batzoglou 2006, Notredame 2007). In an evolutionary context, MSAs are assembled under the assumption that all aligned sequences have diverged from a common ancestor through a series of mutational processes including residue substitution, subsequence insertion and subsequence deletion (collectively insertions and deletions are termed indels). Complete MSAs of sequences under these assumptions are sometimes called global MSAs; for a review of global MSA methods see Notredame 2007. The increasing ubiquity of whole genome sequences has led to an interest in MSAs for complete genomes, including all sequences: genes, promoters, repetitive regions, etc. Termed whole genome alignment (WGA), this requires the aligner to additionally consider genome rearrangements, such as inversions, translocations, chromosome fusions, chromosome fissions and reciprocal translocations. Some tools for WGA are also capable of modelling unbalanced rearrangements that lead to copy number change, such as tandem and segmental duplications, or even retrotransposition mediated duplications (Blanchette et al. 2004, Miller et al. 2007, Paten et al. 2008, Angiuoli and Salzberg 2011, Paten et al. 2011).

WGA methods have been critical to understanding the selective forces acting across genomes, allowing evolutionary analysis of many potential functional elements (ENCODE Project Consortium 2012), and in particular, the identification of conserved non-coding functional elements (Drosophila 12 Genomes Consortium 2007, Lindblad-Toh et al. 2011), including cis-regulatory elements (Kellis et al. 2003), enhancers and noncoding RNAs.

The lack of accepted gold standard reference alignments has made it hard to objectively assess the relative merits of WGA methods. The Alignathon competition presented here is an attempt to redress that imbalance. Previous evaluations of MSAs can be split into roughly four types: those using simulation, those using expert information, those using direct statistical assessments and finally those that assess how well an alignment functions as a component of a downstream analysis. We briefly describe and review these approaches; for a more comprehensive review of some of these methods see Lantorno et al. 2014.

In simulation evaluations a set of sequences and an alignment is generated using a model of evolution. Alignments are created from the simulated sequences and the resulting predictions are compared to the “true” simulated alignment. There are two basic types of simulators for DNA sequence evolution, coalescent simulators and non-coalescent forward-time simulators (Carvajal-Rodríguez, 2010). Forward-time simulators typically model the evolutionary processes in a step-by-step manner and have the drawback that they may spend a great deal of time generating samples or events that are eventually deleted or otherwise go unobserved at the end of the simulation. Coalescent simulators make use of coalescent theory to generate only the observed samples (Hein et al. 2005, Wakeley 2008). While useful for modelling populations, coalescent simulators can not yet efficiently model general sequence evolution, and as a result most simulators for MSA use forward-time approaches.

There are several forward-time simulators useful for assessing global MSA tools. One of the first was ROSE, which is capable of simulating indels, substitutions, and sequence motifs (Stoye et al. 1997, 1998). In assessing the TBA alignment program, Blanchette et al. (2004) added general and lineage-specific transposons and increased rates of substitution in CpG dinucleotides to ROSE to create SIMALI. DAWG is a popular simulator that, similarly to SIMALI, was developed to overcome limitations of ROSE, and features a more general continuous time substitution model (general reversible), and models indel lengths more robustly (Cartwright 2005). Numerous other simulators have been developed for modelling specific mutations or scenarios, e.g. EvolSimulator models lateral gene transfer, and there are several protein-only simulators (Ovcharenko et al. 2005, Beiko and Charlebois 2007, Pang et al. 2005, Nuin et al. 2006, Strope et al. 2007). Varadarajan et al. (2008) have created perhaps the most complex and theoretically elegant set of simulator tools for global MSA currently available — their tools allow for models to be specified by evolutionary transducers and phylogenetic context-free grammars, and can be combined to model numerous features of sequence evolution.

The simulation options for assessing WGA have until recently been absent, essentially because to do so requires modelling both low level sequence evolution and higher level genome rearrangements — a formidable challenge given the large and complex parameter space that potentially encompasses all aspects of genome evolution. The sgEvolver simulator (Darling et al. 2004, 2010) is used to generate simulated genome alignments, though it lacks an explicit model for sequence translocation or mobile element evolution. The ALF simulator (Dalquen et al. 2012) is another choice for WGA, promising to comprehensively model gene and neutral DNA evolution. However, for this study we used the EVOLVER software which can simulate full-sized, multi-chromosome genome evolution in forward time (Edgar et al. 2009). EVOLVER models an explicitly haploid genome and lacks a population model; its framework and expert-curated extensive parameter set are intended to produce “reference-like” genomes, i.e., haploid genomes that represent an amalgam of several individuals. EVOLVER models DNA sequence evolution with sequence annotations; a gene model; a base-level constraint model; chromosome evolution, including inter- and intra- chromosome rearrangements; tandem and segmental duplications; and mobile element insertions, movements, and evolution.

An alternative approach to assessing MSA uses expert biological information not available to the aligner. While interpreting the results of simulations is made difficult by the uncertainty to which they approximate reality, the clear advantage of using expert information is that it can be used to assess alignments of actual biological sequences. For protein alignment there are several popular benchmarks that provide either reference structural alignments or expertly curated alignments (see Blackshields et al. 2007 for review), e.g. BAliBASE (Thompson et al. 2005) and SABmark (Walle et al. 2005). For RNA, established datasets based on structural comparisons include Bralibase (Wilm & Steger 2006) and BraliDARTS (Kemena et al. 2013). Nontranscribed DNA alignments are, however, much harder to assess since one lacks an objective external criterion to assemble objective gold standard references (Kemena and Notredame, 2009). This explains why untranslated DNA alignments are usually evaluated using more ad hoc assessment procedures.

Margulies et al. (2007) assessed WGAs of the ENCODE pilot project regions. To assess misalignment of non-coding regions, the checked for alignments between Alu sequences, a primate specific transposon family, to non-primate sequences. Paten et al. (2008) looked at ancestral repeat alignments within WGAs to see how closely each sub-alignment of a set of ancient repeat homologs recapitulated the known consensus of the ancient repeat. They also addressed the relative amounts of human sex chromosome X aligned to other chromosomes, which, because X is hemizygous in male, is observed to be less prevalently rearranged with the autosomes. The main strength of these procedures is to provide an objective evolutionary context when evaluating the alignment. They are nonetheless weakened by their explicit dependence on the sequence alignment procedures used to determine ancestral repeat relationships.

The difficulty with relying upon expert information is that it may address only a small fraction of the alignment (e.g. coding exons or ancient repeats), and have unknown variance, generality and discriminative power. An alternative approach is to address alignments by statistical measures. For global MSA the Heads or Tails (HoT) measure assesses concordance between the alignment of sequences in forward and reverse complement orientations, reasoning that portions of the alignment that are identical between the forward and reverse alignments are more likely reasonable (Landan and Graur 2008). The GUIDANCE program tests global progressive alignments (those constructed using a guide tree), by creating an ensemble of alignments based upon variations of a base guide tree and establishing regions that are consistent between alignments (Penn et al. 2010a, 2010b). The T-Coffee CORE index (Notredame & Abergel 2003) uses a similar principle and relies on consistency across all alternative pairwise alignments to assess the reliability of individual positions. Clearly, while these metrics may be useful for reasoning about regional reliability they are easily gamed (and possibly inadvertently so, Wise 2009), and thus inappropriate as overall alignment benchmarking methods. A powerful approach was introduced by the StatSigMA-w tool (Chen & Tompa 2010), which assesses the likelihood that sets of homologous residues arose from a given phylogenetic process. If this tool could be scaled efficiently and made to work on arbitrary input it might be a good tool for future work on WGA assessment. For this work we expand on the probabilistic sampling-based alignment reliability (PSAR, Kim & Ma, 2011) method, which samples pairwise suboptimal alignments to assess reliability of MSAs.

Statistical measures are attractive because they can be used with the complete alignments of real sequences. However, without a gold standard to compare against, they are only an uncertain proxy to a true assessment of accuracy. The final category of common assessment methods addresses how well a program generates alignments for a given computational task. This is typically the assessment made by a biologist in choosing an alignment program — how well does it perform in practice, according to intuition or analysis? Unfortunately, these assessments, often being one-offs, rarely make it into the literature, and are difficult if not impossible to generalize from, because the assessment is made for the purposes of the analysis. Notably for WGAs, Bradley et al. assessed how much alignment methods influenced de-novo ncRNA predictions and Margulies et al. analysed the effect of different WGAs on the prediction of conserved elements (Bradley et al. 2009, Margulies et al. 2007).

The Alignathon is a natural intellectual successor to the Assemblathon collaborative competitions (Earl et al. 2011, Bradnam et al. 2013). The starting point of the Alignathon is to assume that the problem of genome assembly is largely a solved problem. Though we admit this is currently a dubious assumption, it appears that the problem of genome assembly will shrink in size in the coming years as new sequencing technologies become available and existing assembly software is perfected to take advantage of more numerous, longer and less error-prone reads. With this future as a starting point, the question a biologist faces changes from being one of “how do I best assemble the genome of my favorite species?” to a higher level question of “how is my favorite species related to the pantheon of other sequenced species?” Such a question is answered through a WGA. If organized community efforts to sequence large numbers of genomes, such as the Genome 10K for vertebrates and insect 5K for insects, are to maximally fulfill their promise by revealing and refining the evolutionary history of all of their species then it is vital that we have the best possible methods for WGA (Genome 10K Community of Scientists 2009, i5K Consortium 2013).

## Results

Of the four discussed strategies to assess alignments we pursue two: simulations and statistical assessment. We do not discount the importance of other methods of assessment, but make a case that we can usefully discriminate the submissions’ performance using those given. We begin by describing the Alignathon datasets in detail before describing the submissions we received, how the submissions were processed, the evaluations that were performed and the performance of the submissions.

### Datasets

The Alignathon used three test sets. Two of the test sets were created by way of forward-time simulation, using the EVOLVER tool, starting from a ∼1/20th scale mammalian genome, a genome size of 120 megabases (Mb), based upon a subset of hg19/GRCh37 (chromosomes 20, 21 and 22; see Methods). The first simulated dataset models a great ape phylogeny consisting of genomes with the same evolutionary relationships as humans, chimpanzees, gorillas and orangutans (Fig. 1). The second simulated dataset is based upon a mammalian phylogeny consisting of genomes with the same evolutionary relationships as humans, mice, rats, cows and dogs (Fig. 1). On a gross level, the summary statistics of the two simulated datasets are shown in Tables 1 and S1. After an initial burn-in phase to shuffle the original input sequences, and so prevent any entrant from using the original source sequence to their advantage (see Methods), the primate phylogeny contained, amongst other changes, one chromosomal fusion and over three million substitutions in the lineage from the MRCA to the simulated human. The mammal phylogeny contained, amongst other changes, two chromosomal splits, one fusion and over 27 million substitutions in the lineage from the MRCA to the simulated human.

**Figure 1.**
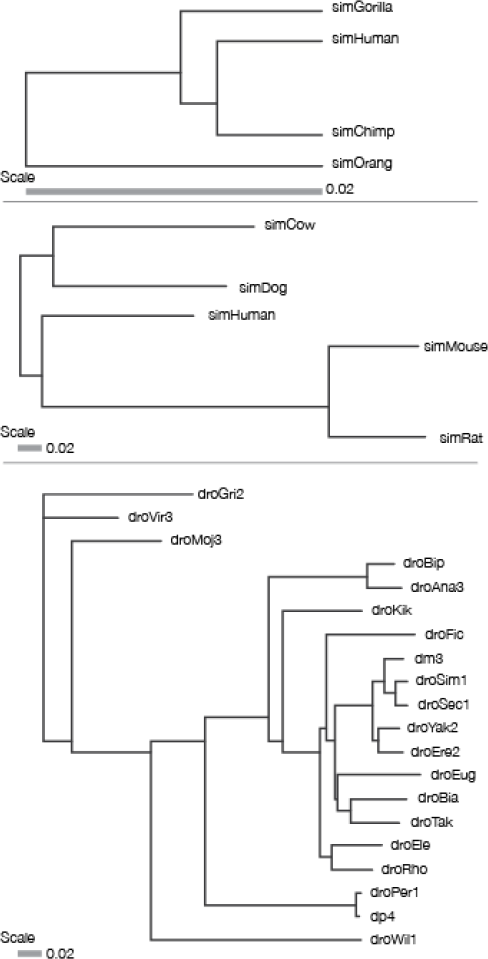
The phylogenies of the three test sets: primate simulation, mammal simulation and fly real data set. Branch lengths are in units of neutral substitutions per site.

**Table 1.**
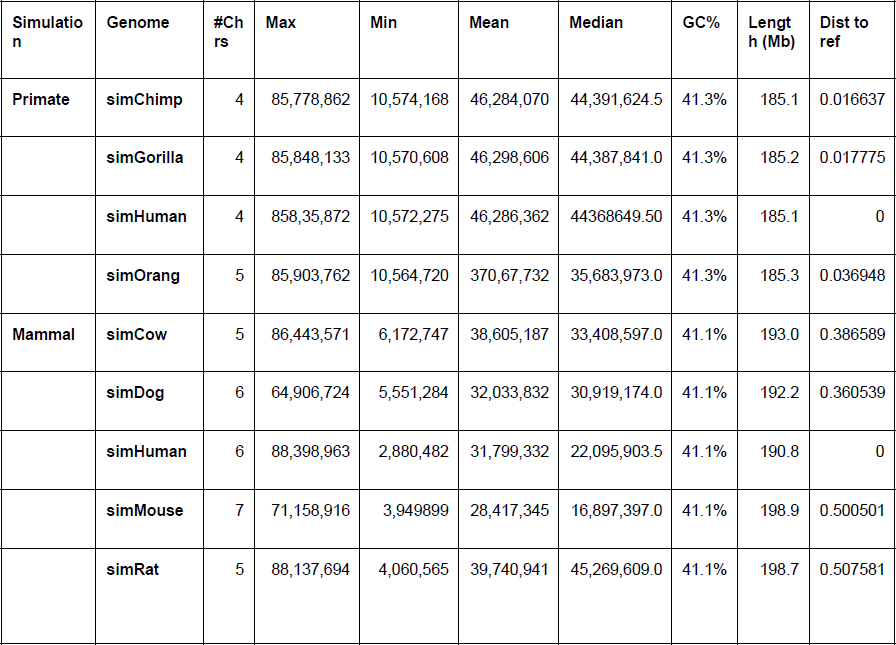
Summary statistics for the simulated genomes. Rows are leaf genomes generatec by simulation. Columns are different metrics: #Chrs is the number of chromosomes; Max is the longest chromosome; Min is the shortest chromosome; Mean is the average length of a chromosome; median is the median chromosome length; GC% is the percent GC composition of the genome; Length is the total length of the genome in megabases (Mb); Dist to ref is the phylogenetic distance from the leaf to the reference species (named simHuman in both simulations).

| Simulation | Genome | #Chrs | Max | Min | Mean | Median | GC% | Length (Mb) | Dist to ref |
| --- | --- | --- | --- | --- | --- | --- | --- | --- | --- |
| **Primate** | **simChimp** | 4 | 85,778,862 | 10,574,168 | 46,284,070 | 44,391,624.5 | 41.3% | 185.1 | 0.016637 |
|  | **simGorilla** | 4 | 85,848,133 | 10,570,608 | 46,298,606 | 44,387,841.0 | 41.3% | 185.2 | 0.017775 |
|  | **simHuman** | 4 | 858,35,872 | 10,572,275 | 46,286,362 | 44368649.50 | 41.3% | 185.1 | 0 |
|  | **simOrang** | 5 | 85,903,762 | 10,564,720 | 370,67,732 | 35,683,973.0 | 41.3% | 185.3 | 0.036948 |
| **Mammal** | **simCow** | 5 | 86,443,571 | 6,172,747 | 38,605,187 | 33,408,597.0 | 41.1% | 193.0 | 0.386589 |
|  | **simDog** | 6 | 64,906,724 | 5,551,284 | 32,033,832 | 30,919,174.0 | 41.1% | 192.2 | 0.360539 |
|  | **simHuman** | 6 | 88,398,963 | 2,880,482 | 31,799,332 | 22,095,903.5 | 41.1% | 190.8 | 0 |
|  | **simMouse** | 7 | 71,158,916 | 3,949899 | 28,417,345 | 16,897,397.0 | 41.1% | 198.9 | 0.500501 |
|  | **simRat** | 5 | 88,137,694 | 4,060,565 | 39,740,941 | 45,269,609.0 | 41.1% | 198.7 | 0.507581 |

Recognizing the limitations of simulations, our third test set consisted of 20 real fly genomes (Fig. 1). The fly genomes were available in various states of completion from near-finished in the case of *Drosophila melanogaster* (dm3 assembly, chromosome sequences) to fragmentary in the case of *D. rhopaloa* (droRho assembly, 34,000 contigs) (Table 2). The fly genomes, while more numerous than the simulated data, are still in the same general size range of the simulated genomes at ∼200 Mb. The decision to use datasets at this size scale, rather than at the scale of a larger vertebrate genome, was balanced, just as in the first Assemblathon, by the desire to attract the largest possible number of entrants while still creating a reasonable challenge.

**Table 2.**
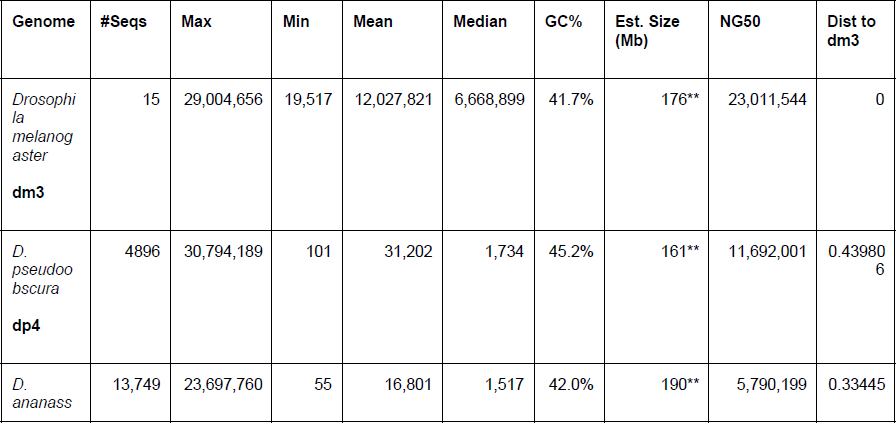
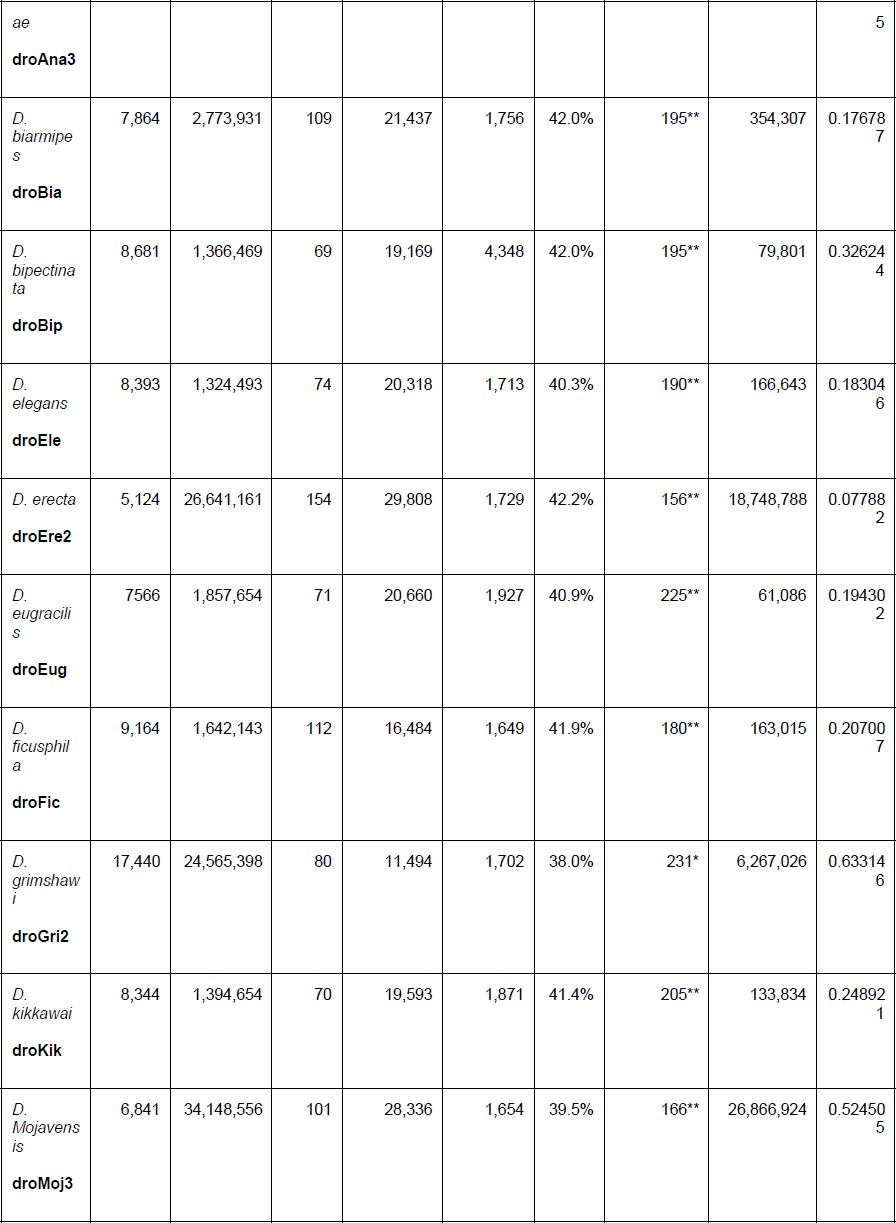
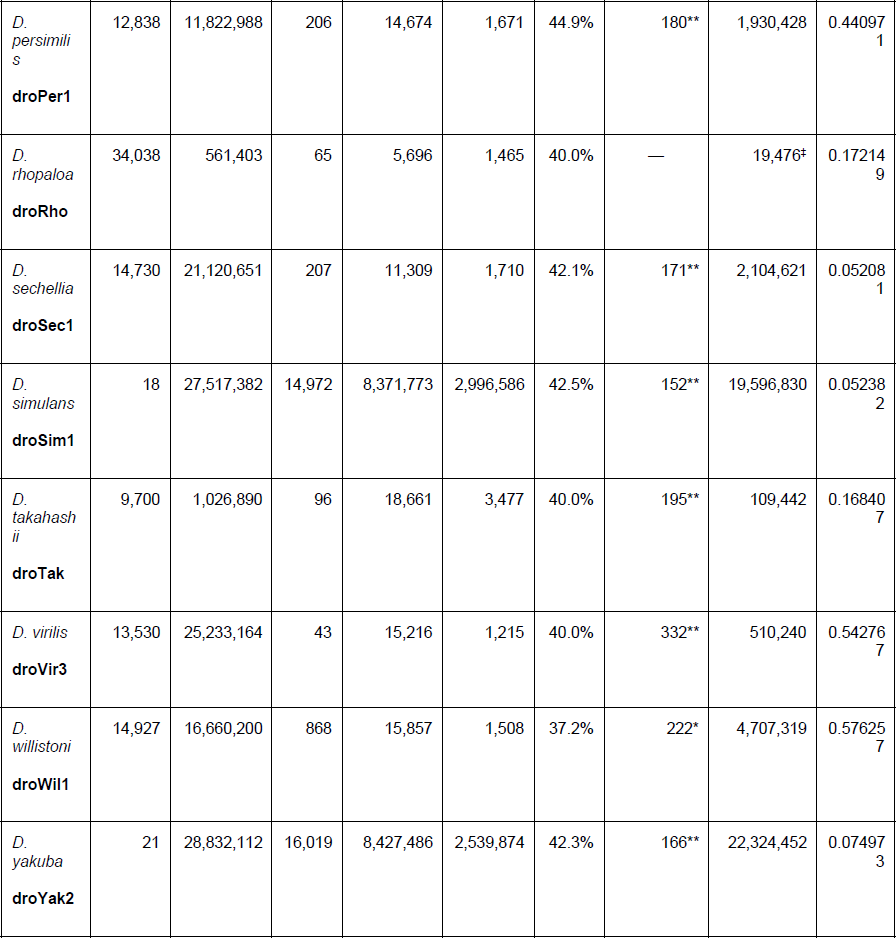
Summary statistics for the fly genomes. For each species provided to participants the following information is shown: number of sequences that comprise the genome; the maximum length of a sequence; the minimum length of a sequence; the mean length of all sequences; the median length of all sequences; the percent GC content of the genome; the estimated size in megabases; the NG50 value of the genome (Earl et al. 2011); the phylogenetic distance to *D. melanogaster* (dm3). ^*^Genome size estimates from Bosco et al. 2007. ^**^Genome size estimates from Gregory & Johnston 2008. ^‡^N50 instead of NG50 value due to lack of genome size estimate.

### Competition Organization and Submissions

The initial datasets were released in December 2011, and teams were given until February 2012 to submit their entries. As in the Assemblathons, none of the teams had access to the datasets until their initial release. The Alignathon received 35 submissions, 13 for the primate simulation, 13 for the mammal simulation and nine for the fly dataset (Tables 3, S2, S3 and S4). Unfortunately, not all algorithms / pipelines were run for all test sets. Participants cited limitations of the methods applied (e.g. inability to handle the scale of the fly dataset), and of resources (time, person-hours, funding, etc.) as reasons for not participating in all datasets.

**Table 3.** Submissions to the Alignathon. Each row shows a tool with: the name of the submission as used in this paper; the names of the submitters; the number of submissions from the tool for the primate data set; the number of submissions from the tool for the mammal data set; the number of submissions from the tool for the fly data set.

| Submission | Tools | Submitter(s) | Primate Simulation | Mammal Simulation | Fly data set (real) |
| --- | --- | --- | --- | --- | --- |
| AutoMz | AutoMz | Minmei Hau | 1 | 1 | 1 |
| Cactus | Cactus | Benedict Paten, Glenn Hickey | 1 | 1 | 1 |
| EPO | Enredo, Pecan, Ortheus | Stephen Fitzgerald, Kathryn Beal, Javier Herrero | — | 1 | — |
| Pecan | Mercator, Pecan | Stephen Fitzgerald, Kathryn Beal, Javier Herrero | 1 | 1 | — |
| GenomeMatcher | Genome Matcher | Igor Seledtsov, Vladimir Molodtsov, PI: Victor Solovyev | 3 | 3 | 3 |
| Mugsy | Mugsy | Aaron E. Darling | 1 | 1 | — |
| Multiz | Multiz | Brian J. Raney | 1 | 1 | 1 |
| PSAR-Align | PSAR-Align | Jaebum Kim, Jian Ma | 1 | 1 | — |
| progressiveMauve | progressiveMauve | Aaron E. Darling | 1 | — | — |
| Robusta | Robusta | Carsten Kemena, Jia-Ming Chang, Ionas Erb, Cedric Notredame | 1 | 1 | 2 |
| TBA | TBA | Minmei Hau | 1 | 1 | 1 |
| Vista-Lagan | Vista-Lagan | Alexander Poliakov, Michael Brudno, Inna Dubchak | 1 | 1 | — |
| **TOTAL** |  |  | **13** | **13** | **9** |

### Genomewide Comparison to Simulated Genome Alignments

All submissions were received in multiple alignment format (MAF) (http://genome.ucsc.edu/FAQ/FAQformat.html#format5). A suite of MAF comparison tools was developed for the project including a comparator tool, so-called because it compares two alignment files. We call the set of aligned pairs of residues within an alignment its *alignment relation*. The comparator tool works by taking two input MAF files, A and B, and comparing their alignment relations. For the simulated datasets, if A is the predicted alignment created by a tool and B is the simulated “truth”, then the ratio of the number of pairs in the intersection of A and B to the number of pairs in A is the *precision* of the prediction. Conversely, the ratio of the number of pairs in the intersection of A and B to the number of pairs in B is the *recall* of the prediction. One standard method for combining precision and recall into a single value is the balanced F-score, which is simply the harmonic mean of precision and recall:

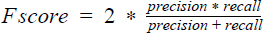

(Beitzel 2006).

The cardinality of the alignment relation of the considered WGAs is exceedingly large, e.g. approximately 1.7 billion pairs for the simulated mammalian alignment. This made complete comparison of the alignment relations impractical. Instead, for each pair of MAFs compared we sampled a subset of the alignment relation of one and checked if any or all elements of the subset were present in the alignment relation of the other, allowing us to affordably estimate the given metrics. By using very large sample sizes (10 million pairs sampled for each direction of a MAF pair comparison) we achieved very high accuracy (data not shown). In the methods we discuss how we sampled aligned pairs in an unbiased and efficient fashion.

For the simulated datasets we performed analyses both with respect to the entire genome and to areas of the genome subsetted by annotation type (genic, neutral and repetitive). Results are shown in Fig. 2 and Tables S5, S6, S7 and S8. We find that many of the submissions were able to align the primate dataset with both relatively high recall and precision, and, with the exception of the GenomeMatch submissions, which had lower values in the repetitive regions, performance was consistently high across annotation types. In particular, the top eight submissions differed by only 0.007 in F-score, and all had recall and precision above 0.98.

**Figure 2.**
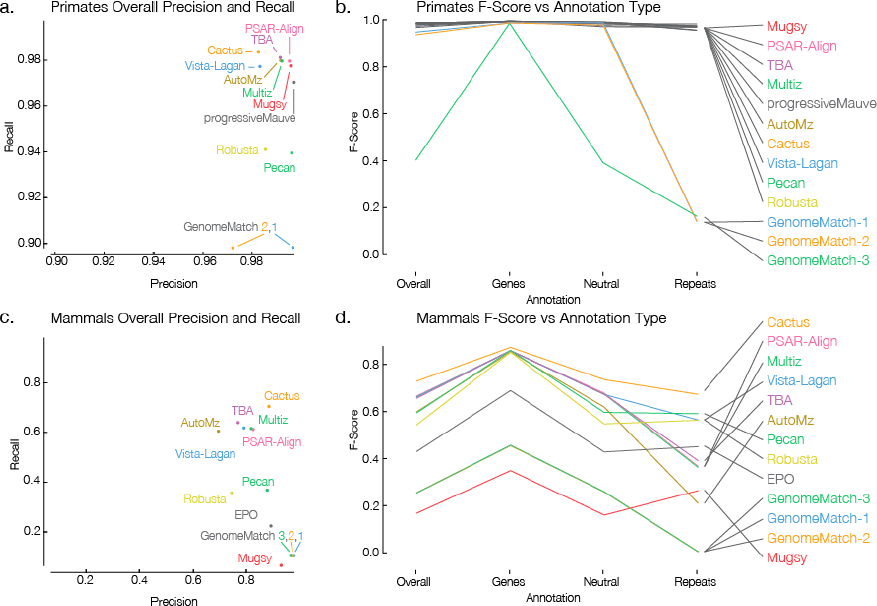
Simulated primate and mammal F-score results. Recall as a function of precision is shown for primates in A and mammals in C. GenomeMatch-3 is omitted from plot A as both of its values are low (see its overall F-score in B). B shows the primate F-score results isolated to different annotation types: overall, genes, neutral and repetitive regions. D shows the mammal version of C. Legends for B and D are ordered as in the overall category and this order is maintained in the genes and neutral annotations.

For the mammal simulations we found a much wider spread of results, both between aligners and within different annotation classes. The strongest submission, Cactus, had an F-score 0.081 points higher than its nearest competitor. Looking at the mammal results by annotation type, generally (and predictably) submissions performed the best in genic regions where simulated selection was presumably highest, performance was intermediate in neutral regions, and submissions generally performed most poorly in repetitive regions. Generally submissions retained their ranking across annotation regions, that is to say that the submissions ranked 1 and 2 in overall were also ranked 1 and 2 in genic regions. However, this trend did not hold for repetitive regions, and surprisingly several submissions performed better in repetitive regions than in the neutral regions (Mugsy, Pecan, EPO, Robusta).

There are some caveats to consider in interpreting these results. For simplicity of interpretation, homologies that predated the MRCA of the extant genomes were not included in the “true” simulated alignments, therefore some ancient homologies captured by the aligners may be considered false in the comparison. Additionally, EVOLVER does not track the alignments it generates when creating simulated transposon insertions, thus two highly similar transposon copies from separate insertion events are not considered homologous in the true alignment. For these reasons, the precision values may be considered a lower bound that may, for some purposes (detecting ancient and transposon mediated alignments) underestimate the accuracy of the alignments to some extent. In addition to these methodological considerations, similarly parameterized EVOLVER simulations were used to benchmark the Cactus aligner in its initial publication, though at approximately 1/250th the scale used here (Paten et al. 2011). It is therefore difficult to know if its increase in relative performance is partially an artifact of training Cactus to the EVOLVER evolutionary model. The GenomeMatch submissions did not attempt to align the repeat-masked sequences, which may explain their poor performance in repeats, and therefore overall, but stronger relative performance in genes and, to some extent, neutral regions.

As phylogenetic distance between species grows the number of unobserved mutation events increases and the alignment problem naturally becomes more difficult (Wong et al. 2008, Landan & Graur 2008, Holmes & Durbin 1998). To see this, we stratified the results by phylogenetic distance (path length between leaves in the simulated phylogenies) between all pairs of species; see Fig. 3. Longer distances are indeed observed to lead to lower precision and recall values, and therefore lower F-score values. For reference based aligners, which use one species as a reference (here simHuman), there is a clear dip in performance for non-reference pairs (pairs not including the reference sequence). This is especially prevalent in Fig. 3.B for the PSAR-Align submission, which used the Multiz program, and the Multiz and AutoMz submissions, which rely upon the Multiz program. The TBA program, which was developed by the Multiz authors to overcome this issue, does not suffer from the problem.

**Figure 3.**
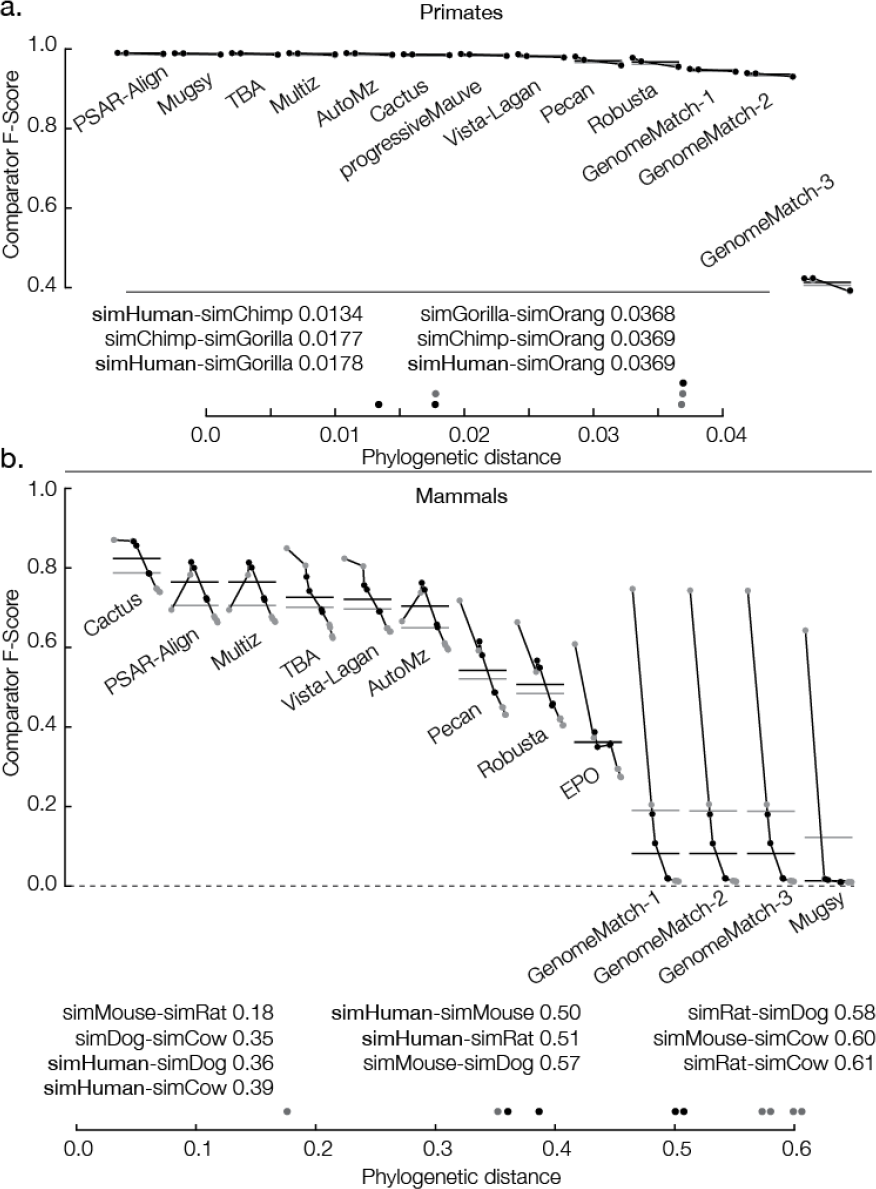
Primate and Mammal simulation F-score results stratified by phylogenetic distance. For each subplot the vertical axis shows the F-score and the horizontal axis shows 13 individual submissions ordered from left to right (descending) by average overall F-score. Horizontal gray lines show the overall F-score of the submission taking into account all sequence pairs. Horizontal black lines show the overall F-score of the submission taking into account only sequences pairs including the reference. Submissions are comprised of points connected by a line where the points are in ascending order of phylogenetic distance (all possible pairs are shown).

### Evaluating Genome Alignments in the Absence of a True Alignment

We sought statistical measures that we could compare to the simulation results and use to assess the fly dataset. We attempted to use the StatSigMA-w tool (Chen & Tompa 2010), but the version provided by the authors does not work with arbitrary input data. Instead, we used the recently developed PSAR tool (Kim & Ma 2011). PSAR assesses an alignment by removing a sequence, sampling suboptimal alignments between the removed sequence and the remaining alignment using the forward algorithm with a pair-HMM (Durbin et al. 1998), and then checks to see how well the newly sampled alignments match the original alignment. By repeatedly performing this sampling with every possible sequence PSAR is able to calculate alignment reliability scores for every pair of matched residues in the alignment. Each score is similar to a posterior probability that a given pair of residues in the input alignment are aligned (Durbin et al. 1998), i.e. it can be thought of as a proxy to a local measure of accuracy, which factors in the edit matrix surrounding the pair of aligned residues. We define the *PSAR-precision* for a pair of genomes as the average of the PSAR pair scores for the matched pairs of residues in the alignment including both genomes.

To deal with its limited alignment model — which is appropriate for global MSA, allowing only substitutions, insertions and deletions — and to make it computationally feasible to assess the alignments, for each of the datasets separately we ran PSAR on five half-megabase subregions (see Methods), using sampled intervals of a chosen reference genome to pick the subregions (for the flies *D. melanogaster*, dm3, and for the simulations simHuman), and manipulating the alignments to make them appropriate for PSAR (e.g. removing duplications, see Methods and below). The overall *PSAR-precision* for the complete alignment is the average of PSAR-precisions for genome pairs including the reference. The PSAR-precision scores are analogous to the precision measures calculated from the simulations, because they estimate the expected number of pairs in the alignment that are correctly aligned.

The advantages and limitations of using PSAR for assessing WGAs are important to note. Firstly, any rearrangements within a sampled region of the non-reference species are invisible to the PSAR method, and so will not disrupt the PSAR score. Secondly, the submitted alignments (and biological reality), may contain duplications within a subinterval, and these are simply removed by a principled, but heuristic methodology (see Methods). Thirdly, we break up the PSAR computation of a region into smaller “chunks” (see Methods). The chunking is likely to have somewhat affected the scores, though in our experience this proved a very minor effect, as the number of breaks between chunks is very small compared to the number of columns in the alignment. Fourthly, to make the scores comparable between different alignments, we look only at pairs including the reference sequence, which is a shared constant — and thus ignore non-reference pairs. Finally, and perhaps most importantly, PSAR is estimating accuracy based upon alignments sampled from a singly parameterised pair-HMM, its objective function is therefore somewhat biased towards similarly parameterised models.

To complement our proxy to precision we used a simple proxy to recall: coverage. For a pair of genomes A and B, the proportion of residues in A aligned to a residue from B is the *coverage of B on A*. The *overall coverage* (where we drop the ‘overall’ when it is clear from the context) is the average of coverages for all pairs of distinct species. Hypothesizing that PSAR-precision can used to approximate precision and that coverage can be used as an estimate of recall, the natural statistical analog to F-score is the harmonic mean of PSAR-precision and coverage, which we call the *pseudo F-score*.

To see how consistent our statistical measures were to the measures derived from the simulations we calculated them for the user-generated simulated primate and mammal alignments (Fig. 4 and Tables S5 and S6). In the simulated primates all the values were uniformly high, and hence saturated. However, looking at the simulated mammals we find a very good, linear correlation between recall and coverage (*r*^2^ = 0.984), but no linear correlation between PSAR-precision and precision. In particular PSAR reports relatively consistent, high scores for all the different alignment programs, suggesting that at a local, residue level on aggregate the alignments look equivalently reasonable (given the caveats mentioned above) between alignment programs (i.e. have about the same number of suboptimal alternatives). It should be noted that PSAr uses the same pair-HMM alignment model as used by the PSAR-Align team in generating their alignments. We might expect therefore that the PSAR-Align alignments would be judged most accurate, though we actually found a number of other programs earned equivalently high results. Despite the lack of linear correlation between precision and PSAR-precision, we find that, because of the excellent recall and coverage correlation, the F-score and pseudo F-score results linearly correlate strongly (*r*^2^ = 0.975 in simulated mammals). This appears to be because the more limiting factor in many of the alignments performance was not precision, but rather a lack of relative recall/coverage, something particularly affecting the GenomeMatch, Mugsy, and, to a lesser extent, Pecan, EPO and Robusta submissions.

**Figure 4.**
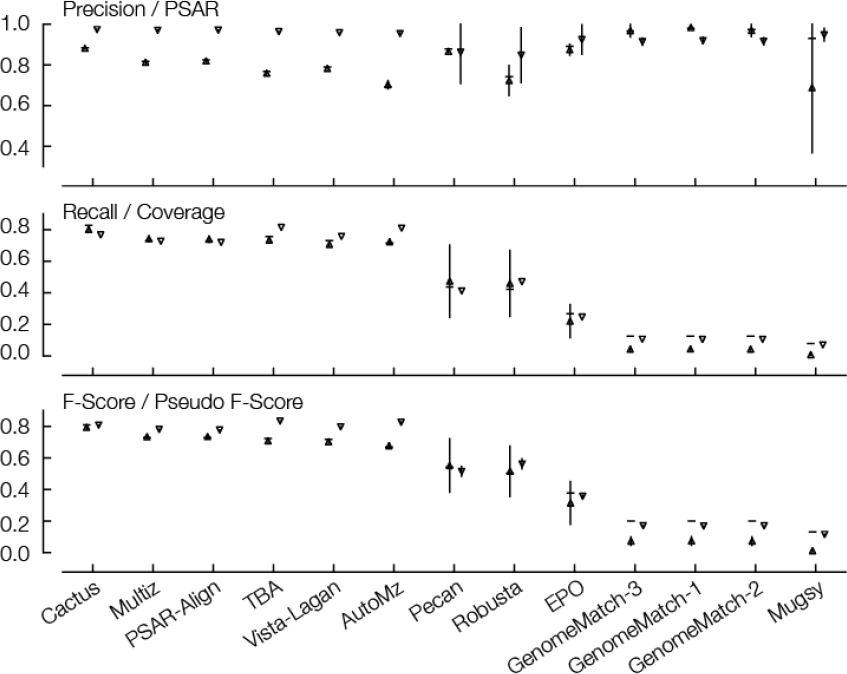
Simulated mammal results comparing simulation values to statistical values. Shown is precision and PSAR precision; recall and coverage; F-score and pseudo F-score. Each column represents the results of one submission, columns are in descending order of overall (full genome) F-score value. The horizontal line is, respectively, the overall precision, recall or F-score value; the up-pointing triangle with a vertical line is the regional precision, recall or F-score mean value, +/− the regional standard deviation; the down-pointing triangle with a vertical line is the PSAR-precision, coverage or pseudo F-score mean value, +/− standard deviation for values that were computed using regional sub-alignments.

In addition to looking at the overall correlations, we calculated regional precision and recall values using the regional alignments (see Methods); this checked for any bias created by the use of a set of manipulated regional alignments in calculating the PSAR-precision values (also shown in Fig. 4 and Tables S5 and S6) — and which might explain the poor PSAR-precision vs. precision correlation. Looking at the mammals, we find reasonable linear correlations between regional and overall precision (*r*^2^ = 0.594), recall (*r*^2^ = 0.989) and F-score values (*r*^2^ = 0.992) (Figure S8), suggesting that the alignment manipulations did not bias the results too substantially. Furthermore, we find for most submissions a low variance in these results between different regions, suggesting five regions were likely enough to get a good approximation of the overall results. The only submissions with relatively high regional variation in these results was the Pecan, EPO and Robusta mammalian submissions, which appear to vary more at this fairly large-scale resolution — perhaps indicating the relative coarseness of the synteny maps that were used (see discussion).

Having assessed the relationship between the statistical and simulated measures we are in a position to extend the analysis to the fly dataset in an informed manner; Fig. 5 and Table S9 shows the overall PSAR-precision, coverage and pseudo F-score results. For the teams that submitted alignments for both datasets we see good concordance between the fly and simulated results. Again, we see the difference between the aligners dominated by coverage differences, with uniformly high (all greater 0.97) average PSAR-precision values that mostly lie within the regional standard deviations of one another, with the exception of the GenomeMatch alignments, which have very high PSAR-precision values but relatively low coverage. Perhaps surprisingly given their reference assisted nature, we find that, along with Cactus and TBA, Multiz and AutoMZ had high relative coverage and pseudo F-scores, even when factoring that coverage was calculated over all pairs, not just reference containing pairs. Plotting the pairwise coverages between all pairs of species (Fig. 6) we see that all the programs had higher relative coverage for pairs involving the reference, partially this is an artifact of the structure of the phylogeny (Fig. 1). The reference based aligners (here Multiz and AutoMZ) indeed did have the highest coverage for reference pairs, and the strongest non-reference based aligners by these metrics, TBA and Cactus, showed a smaller separation between reference and non-reference species pairs.

**Figure 5.**
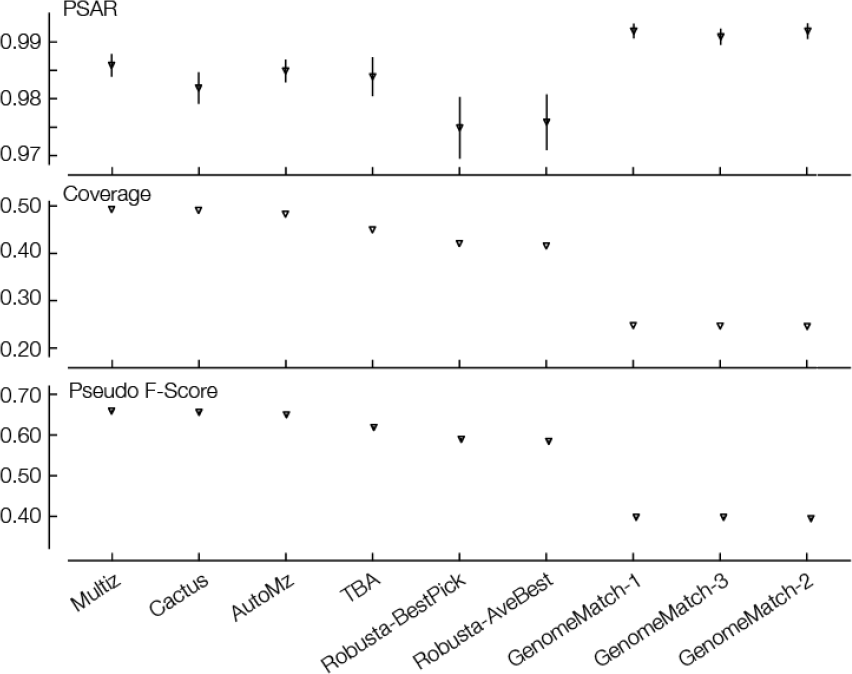
Fly results: values of PSAR-precision; average overall coverage between all pairs; pseudo F-score. Columns are in descending order of mean pseudo F-score value. For each metric, each submission is made up of an orange circle with a vertical line representing the regional mean +/− stdev.

**Figure 6.**
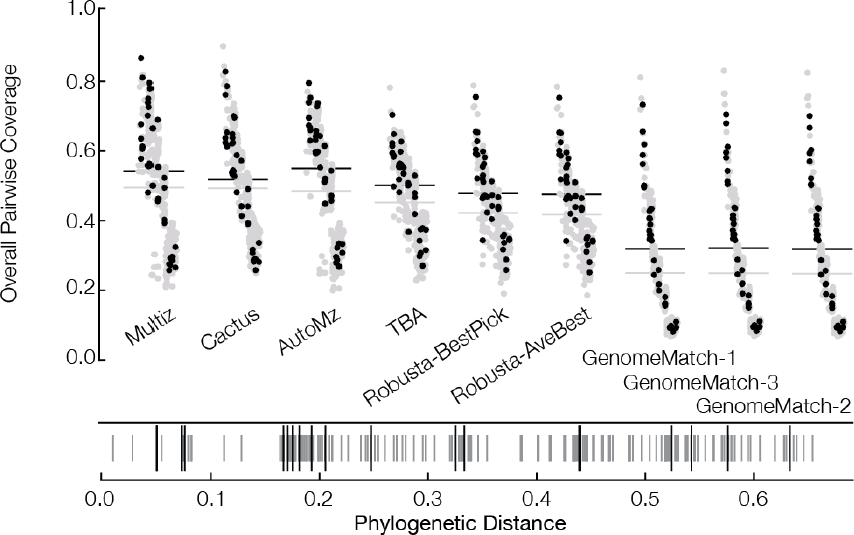
Overall pairwise coverage values in the flies dataset. Submissions are ordered left to right (descending) by overall coverage. Gray points are non-reference pairs, black points contain the reference. The horizontal gray line shows the average coverage of the submission for all points, the horizontal black line shows the average coverage of the submission for just pairs containing the reference. Beneath the pairwise coverage plot is a barcode plot showing the phylogenetic distances of all pairs, shorter gray lines are non-reference pairs and longer black lines are reference containing pairs.

### Visualizing Regional Accuracy

Earlier we demonstrated that the performance of alignments varied regionally according to the simulation of annotation types. Having developed scoring metrics that can be applied across simulated and non-simulated genomes, we are in a position to potentially corroborate this analysis by visualizing how the scores vary across the sampled subregions. To view a complete region at approximately this level of resolution, for each subregion we binned the reference sequence into 1 kb non-overlapping intervals and calculated the F-score (for the simulated datasets) and pseudo F-score for each bin, calculating the score for a bin as if it represented the complete alignment, and for the simulated comparisons, restricting the true alignment to just those pairs involving residues in the reference interval defining the bin. Fig. S1, 7 and 8 visualizes how the scores vary across example regions of, respectively, the simulated primate, mammalian and fly alignments. It is clear that the “best” alignments by these measure differs substantially from the poorest, and that for many submissions there is considerable regional variation. Looking across all the simulated regions, the F-score and pseudo F-score measures correlate reasonably bin-by-bin (Fig. 9, *r*^2^ = 0.671), indicating that pseudo F-score can be used as a reasonable proxy to F-score at this regional level of resolution (e.g. see Fig. S3, the equivalent to Fig. 7 but using pseudo F-score instead of F-score). It should be noted that the correlation is imperfect; in particular it appears that the pseudo F-scores saturate at high values, while the corresponding F-scores still discriminate alignment quality. I.e. pseudo F-scores do not always discriminate between good and very good alignments; this again seems to be an artifact of the lack of good linear correlation between precision and PSAR-precision, rather than between recall and coverage (see Fig S4. and S5).

**Figure 7.**
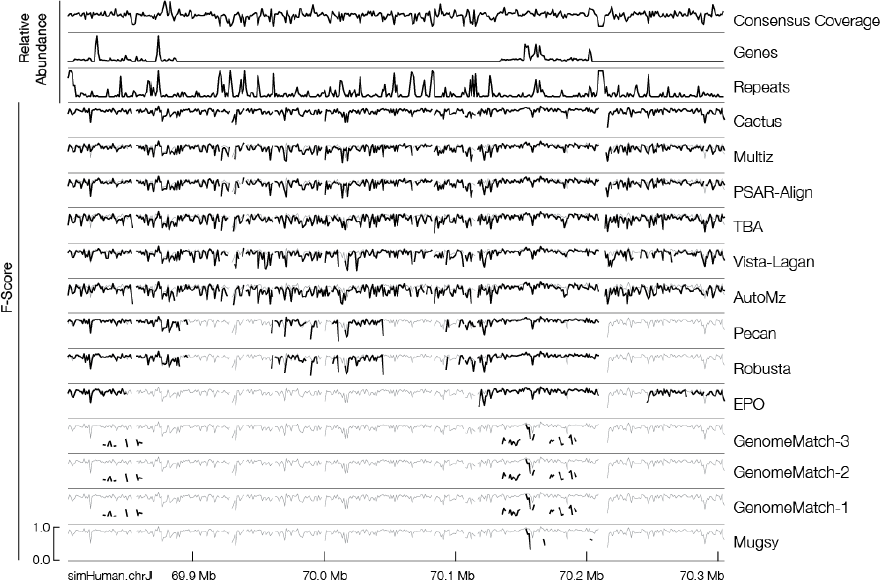
Region 4 of simHuman with respect to simMouse of the regional analysis of the mammal simulation data set. Region 4 is defined as bases 69,805,407 - 70,305,406 of simHuman chromosome J (horizontal axis). Rows are: the relative true coverage of any part of simMouse onto this region of the reference; the relative abundance of genes within the region; the relative abundance of repetitive sequence in the region; submissions in descending order of average F-score. Each submission row shows the F-score of the submission at a given location of the region in black. The vertical axis of each row is the same scale, as labeled in the bottom row. In grey, in the background, is shown the top submission (Cactus). Note that most submissions managed to contain parts of the alignment within the gene regions, though half of the submissions had poor coverage in this region (they lack a black line through most of the plot). The lack of consistent signal from all of the GenomeMatch submissions may be explained by the fact that the submitters excluded repetitive regions from their alignment.

**Figure 8.**
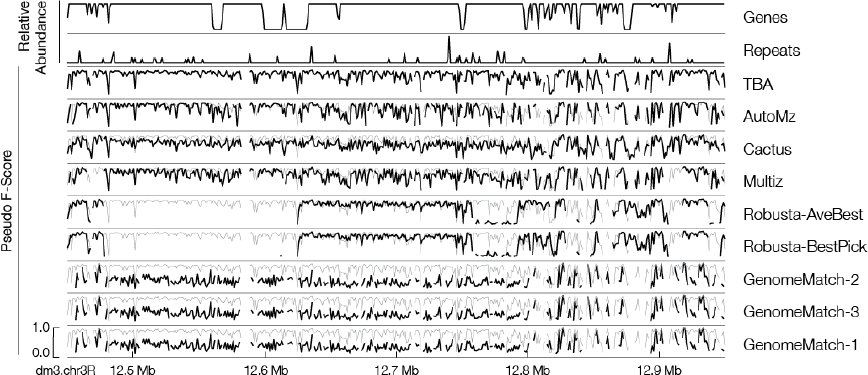
Region 2 of *D. melanogaster* (dm3) with respect to *D. grimshawi* (droGri2) of the regional analysis of the mammal simulation data set. Region 2 is defined as bases 12,450,223 -12,950,222 of dm3 chromosome 3R (horizontal axis). Rows are: the relative abundance of genes within the region; the relative abundance of repetitive sequence in the region; submissions in descending order of average pseudo F-score. Each submission row shows the pseudo F-score of the submission in black. The vertical axis of each row is the same scale, as labeled in the bottom row. In grey, in the background, is shown the pseudo F-score value of the top submission for this region (TBA).

**Figure 9.**
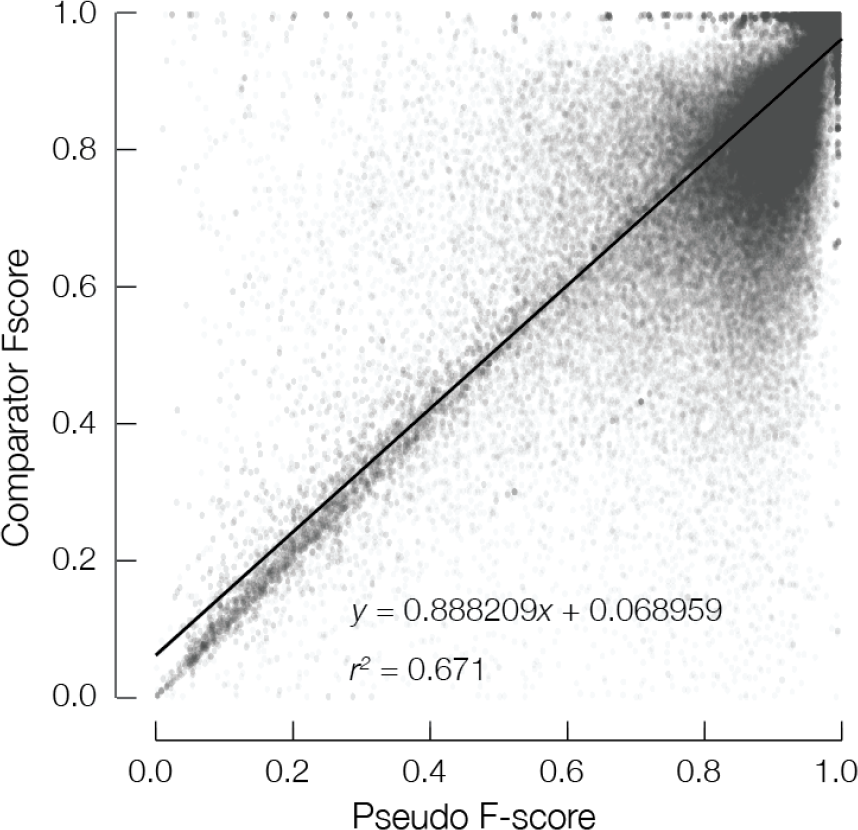
Correlation between PSAR-based pseudo-F-Scores and F-Scores for identical bins with respect to the reference (simHuman for both the simulated Primate and simulated Mammalian data sets) using all pairs of species that include the reference and derived from all submissions. Transparency (*α* = 0.05) is used to represent the data points (*n* = 174,110) such that highly dense areas are darker than others. The strong cloud of data points to the upper right are bins and pairs derived almost entirely from the simulated Primate data set. The overall trend is roughly linear with a coefficient of determination *r*^2^ = 0.671.

### Comparing the Alignment Relations

Several of the pipelines used some of the same underlying programs. To see how these commonalities affected the alignments, for each dataset we calculated the Jaccard distance between the alignment relations of each of the submissions; see Figure 10. As predicted by the earlier analyses, the primate submissions are relatively similar to one another, while the mammalian and fly submissions prove much more divergent. The commonality between submissions is striking, with the same patterns being repeated across the three datasets, and fits well with the programmatic commonalities that the pipelines share. For example, the AutoMZ and Multiz submissions are parameter variants of the Multiz program (Miller et al. 2007), and the PSAR-align and TBA submissions both used elements of the Multiz pipeline. Similarly, the Pecan and Robusta submissions used Mercator (Dewey 2007) to establish the blocks of syntenic sequence and they appear relatively similar. The results indicate that some of the programmatic commonalities between the alignment pipelines are perhaps more important than others, e.g. sharing the same synteny block generator (Mercator or Multiz), had a greater effect on the results than sharing the same synteny block aligner, for example the EPO and Pecan submissions both used the Pecan program (Paten et al. 2008, Paten & Birney 2009) to align sets of syntenic sequences, and the Cactus program (Paten et al. 2011) uses the same pairwise-HMM to generate much of its multiple alignment as Pecan, but these submissions were relatively different from one another.

**Figure 10.**
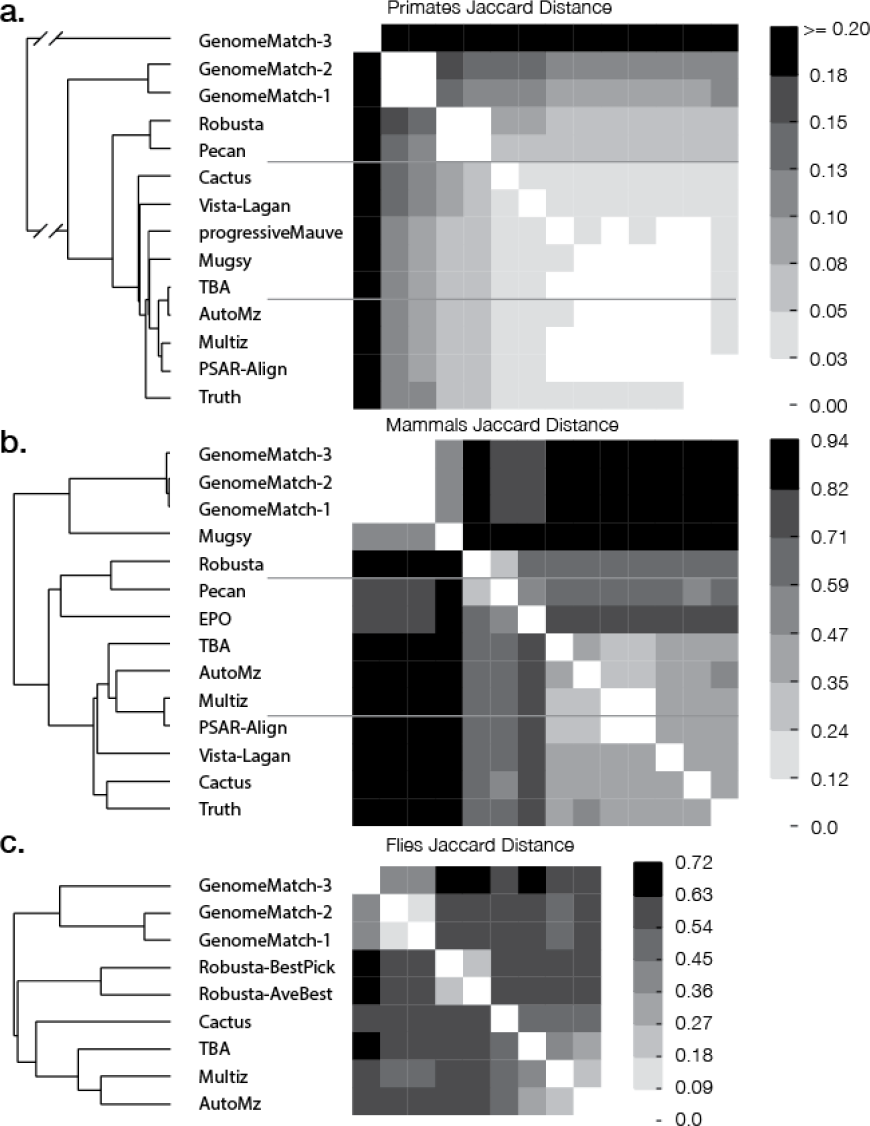
The Jaccard distance (1 - Jaccard similarity coefficient) matrix and accompanying hierarchical clustering (UPGMA) of submissions for each of the three test sets. Higher values indicate that the sets of aligned pairs of two submissions are more dissimilar, lower values indicate similarity.

## Discussion

With the explosion in sequencing delivering ever larger numbers of near complete genome assemblies, WGA is an essential and increasingly important task. We have tested a total of 35 submissions from 12 different pipelines across three different datasets to produce, to our knowledge, the largest and most comprehensive assessment of WGA to date.

The pipelines used to generate the alignments represent those used by genome browsers to generate their WGAs: Vista-Lagan for the Vista Browser (Frazer et al. 2004, Dubchak et al. 2009); Multiz for the UCSC Browser (Miller et al. 2007, Meyer et al. 2013) and Pecan and EPO for the Ensembl Browser (Paten et al. 2008, Paten & Birney 2009, Flicek et al. 2013). In addition, we tested a fairly broad set of standalone WGA tools, including: progressiveMauve (Darling et al. 2010) ; TBA (Blanchette et al. 2004); Cactus (Paten et al. 2011); Mugsy (Angiuoli & Salzberg 2011) ; a meta-WGA tool, Robusta (Notredame 2012), which combines results from multiple standalone tools; and a realignment tool, PSAR-Align, which was used to realign Multiz based alignments in this competition but can in principle refine alignments from any multiple alignment tool. We also tested pairwise WGAs from the GenomeMatch team.

We have demonstrated by simulation that for closely related genome sequences aligners can find the vast majority of homologies accurately — 8 of the 11 pipelines had F-scores above 0.98 in the primate dataset, and only one submission, probably due to exploratory parameters, scored less than 0.9. Conversely, in the simulated mammalian alignments we found a broader distribution of results, with the difference in F-score between the best and worst scoring pipeline being 0.68. In concordance with this, we find that accuracies were substantially higher between more closely related genomes, even in alignments also involving more distantly related genomes — this was apparent both in looking at F-scores in the simulated mammals (Fig. 3), and pseudo F-scores in the flies (Fig. S2).

Testing using both simulations and real data, for those pipelines that submitted results for both fly and simulated mammalian datasets, we find a clear concordance between the rankings. In addition, using the simulated datasets we were able to demonstrate reasonable linear correlations, both overall and regionally, between F-scores and pseudo F-scores. This is important, because it indicates that the high-level aggregate differences we highlight between the submissions can be found by two entirely independent means. Perhaps surprisingly, we did not find a linear correlation between precision and the statistical measure of precision (PSAR-precision) we used, but we did find a very strong correlation between recall and coverage. Importantly, for the submissions we received on both flies and simulated mammals, differences in recall were overall greater than differences in precision, and therefore more critical in determining the observed performance differences.

Although it is encouraging that we found some concordance in the results between the simulated and fly datasets, there are certainly some issues to consider that can likely only be resolved by further work. Firstly, some individual results on the simulations were somewhat surprising. For example, the EPO results had particularly low coverage on the simulations — substantially lower than that pipeline achieved in the genome alignments available from Ensembl (Paten et al. 2008). Unfortunately the EPO pipeline was not applied to the fly dataset, so it is difficult to make a properly controlled comparison, but it is quite possible that, for example, some artifact of the simulation particularly affected their pipeline. Secondly, none of these datasets, intentionally, were of the scale of whole vertebrate genomes, and none contained more than two dozen genomes. It is difficult to predict what increasing the number and scale of the genomes would have done to the comparison. Finally, although these datasets were likely large enough to gain some statistical reproducibility for these assessments and alignments, we do not know how, for example, alternate assessments of the fly dataset or parameters for the simulations would change the picture. Related to this, in general, it is difficult to compare our results to those previously published (see the introduction for a brief overview).

Much previous assessment of WGAs has been made as part of the publication of a novel tool. Naturally, these assessments tend to present results that favour the presented tool. There have been relatively few independent or community organised assessments of WGA pipelines. Notably, as part of the ENCODE pilot project (Margulies et al. 2007), four pipelines were assessed across a substantial number of regions, and Tompa et al. later compared those alignments using the StatSigmaW tool (Chen & Tompa 2010). Unfortunately, direct comparison of such results is complicated by the differences between the methodologies used. For example, Tompa found the Pecan based pipeline to be the most accurate of the four, but here Pecan was run with a different set of preprocessing tools to establish an initial synteny map of the sequences (Mercator and, separately, with Enredo).

### Submission Normalization

An implicit assumption in this assessment is that a WGA represents a partitioning of the residues in the input (extant) genomes into evolutionarily homologous sets. The MAF specification, which alignments were submitted in, has a key ambiguity given our evolutionary assumption that makes apples-to-apples comparisons difficult: a sequence residue may appear in one or more columns of a file. Allowing a residue to appear in multiple columns violates the transitivity of evolutionary homology: E.g. if a residue *x* is aligned to a residue *y* and *y* is aligned to a residue *z*, then *x* and *z* should be aligned, because it is not possible for *x* and *y* to share a common ancestor and *y* and *z* to share a common ancestor while *x* and *z* do not share a common ancestor. Comparing two alignments, one of which is transitively closed, and one of which is not, based only on the aligned pairs contained in a MAF gives an unfair advantage to the non-transitively closed submission. This is because the transitively closed submission must align all residues transitively connected by alignments, which may lower the overall precision of the set of aligned pairs.

Here we assessed a variety of WGAs, some of which are naturally not transitively closed. To see how different the results would be if we were to have enforced transitive closure we created a tool (mafTransitiveClosure) which computes the transitive closure of a MAF (a linear time operation), Fig. 11 shows the results for the simulated datasets. We found that the progressiveMauve, Cactus, Pecan, Robusta, EPO and Mugsy programs produced WGAs that were transitively closed, and therefore unaffected by transformation. As predicted, those submissions that were not initially transitively closed all saw their precision performance decline, in some cases very substantially, and, perhaps surprisingly, no submission saw a significant boost in recall. The GenomeMatch submissions were pairwise, and therefore understandably not transitively closed. Clearly, naively taking the transitive closure of such a pairwise alignment does not necessarily result in a reasonable WGA. The submissions based upon Multiz (Multiz, AutoMZ and PSAR-Align), which are reference based, allow a subsequence of a non-reference genome to align to multiple different subsequences of the reference genome, and are therefore also not naturally transitively closed.

**Figure 11.**
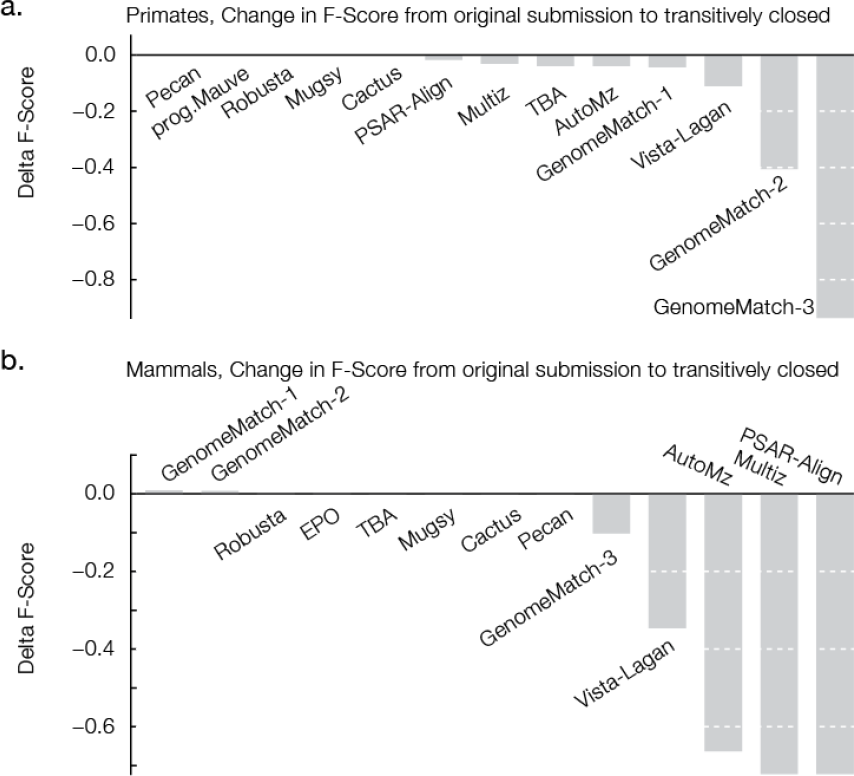
The change in F-Score between original submissions (exactly as provided by participants) and the transitive closure of those same alignments for the simulated primate (A) and simulated mammal (B) data sets. A negative value indicates that following transitive closure the alignment had a lower F-Score.

### Missing Duplications

In theory several of the tools should have been able to align together duplicated regions. To find duplications within the alignments we used a simple metric, for a pair of genomes A and B, the duplicative coverage of B on A is the proportion of residues in A aligned to two or more distinct residues in B. This assessment is made complicated by the lack of transitive closure in some submissions, because a single residue may align to two or more residues in a genome, but in separate columns of the file. To avoid this complexity we assessed the submissions after computing the transitive closure (which made the task computationally significantly easier). To avoid misrepresenting submissions we dropped submissions from the assessment for which the transitive closure adversely (> 0.05 change) affected the F-score or pseudo F-score. Figure S6 shows the results; in short, we find that only Cactus had significantly non-zero duplicative coverages, e.g. just over 3% of all fly genome bases were found to be duplicated, on average, when looking at any other genome. Unfortunately we had only simulation and not fly alignments for other duplication aware tools, such as EPO and Mugsy, so it is hard to draw firm conclusions, but across the pipelines there is likely significant room for WGA tools to improve in detecting duplicative homologies.

### A Code and Data Repository to Reproduce the Simulation Results

One criticism of competitive evaluations like the Alignathon, is that, as one-offs, they do not allow an ongoing determination of best-of-breed. To attempt to mitigate this we have created an easy to evaluate benchmarking pipeline that we hope will continue to be used in future assessments available at http://compbio.soe.ucsc.edu/alignathon/. Unfortunately the PSAR analysis involved using a compute cluster, making it expensive for outside groups to repeat this assessment. However, given a MAF file alignment of one of the simulated datasets, our efficient sampling strategy can be used to make a relatively quick performance assessment. We therefore provide code that makes it trivial to repeat the simulation analysis. The user would download the analysis repository, compile the necessary software, download the requisite data, place their alignment in a specified subdirectory and type “make” in the terminal window to launch the analysis. This approach will hopefully spur future development and assessment upon this resource, though we caution against overtraining to these datasets.

## Conclusion

Robust WGA tools are critical for the future of comparative genomics. We have demonstrated that most of todays tools are generally good at aligning closely related genomes, but that with increasing evolutionary distance WGA’s get progressively less accurate and correspondingly the variation between tools grows substantially. We’ve also highlighted the differences in the types of alignment output (pairwise, reference and referenceless being three obvious subcategories) by WGA tools, and found that finding standard ways to compare between them is not trivial, for example, our computation of the transitive closure proved too disruptive to many alignments to aid useful comparison.

If asked to propose a winner of the competition, it is reasonable to claim that it depends upon the requirements of the user. Users solely interested in a reference based alignment can be reassured that the Multiz based pipelines (Multiz, AutoMZ and here PSAR-align) perform well, but they should be aware that while these pipelines may produce the maximum coverage on a chosen reference species, this focus comes at the expense of a clearly demonstrated bias against alignment between non-reference species. A user looking for a tool to align genic sequences pairwise can reasonably conclude, based upon the evidence of the simulations, that GenomeMatch is an acceptable solution. Users interested in precise alignments, as is often required in phylogenetic inference highly sensitive to errors, and with no need for high recall, might pick progressiveMauve (which displayed excellent precision in the primates).

Given the reasonable uncertainty about the realism of the simulation model and the apparent limited resolution of our statistical metrics, we caution against over-interpretation of the results presented. Perhaps our central conclusion therefore is that more work is needed to confirm or refute our findings, and that generating a wider variety of WGA benchmarks is important for the field to progress on firm ground. Beyond the questions addressed here, we have left a number of big questions unanswered. For example, we have not attempted to determine how tools for WGA compare to methods for other types of MSA, such as protein aligners, or how the quality of the genome assemblies affects WGAs. In summary, we very much hope that the Alignathon will help pave the way for subsequent efforts with more datasets, comparisons, accurate statistical assessments and even broader scope.

## Methods

The project website is available at http://compbio.soe.ucsc.edu/alignathon/. It links to all the datasets, submissions and evaluation code.

### Simulations

As in the Assemblathon 1 project (Earl et al. 2011) simulated genomes were generated using the EVOLVER suite of tools forward-time whole genome evolution simulation tools (Edgar et al. http://www.drive5.com/evolver/). Specific parameter files used to create the simulations are available on the project website. EVOLVER has a model for proteins, genes and base-level evolutionary constraints. EVOLVER uses a two step process for simulating a single forward step in a simulation: the first step is an intra-chromosomal evolution step, and the second an inter-chromosomal step. The intra-chromosomal step allows events such as substitutions, insertions and deletions, duplications, translocations, and inversions, according to rates distributed according to the length of the event. The inter-chromosomal step allows chromosome fusions, fissions, segment copying, segment movement, reciprocal translocations and non-reciprocal translocations. Additionally EVOLVER keeps a separate mobile element library that can insert mobile element DNA into the simulated genome; this library is itself also undergoing simulated evolution. EVOLVER logs all evolutionary events that take place during a cycle and keeps track of the relationships between residues in the parent and child genomes.

EVOLVER as distributed is only capable of performing a single cycle of evolution. In order to run the arbitrary phylogenies necessary for this project we used the evolverSimControl and evolverInfileGeneration tools available at https://github.com/dentearl/evolverSimControl and https://github.com/dentearl/evolerInfileGeneration/. respectively. These extra tools, along with mafJoin https://github.com/dentearl/mafJoin/ were used to construct MAF files containing the entire simulated true evolutionary relationships of all of the genomes: leaves, internal nodes, and the root.

As in the Assemblathon project, we initiated the simulation using a subset of the well annotated human genome, hg19/GRCh37. Complete chromosome sequences for chromosomes 20, 21 and 22 along with annotations for those chromosomes from the UCSC genome browser tracks mgcGenes, knownGene, knownGeneOld5, cpgIslandExt, and ensGene were obtained from the UCSC golden path download site. The tool suite evolverInfileGeneration was used to take the raw data and make it into an EVOLVER infile dataset. This starting dataset was then put through the EVOLVER simulator for a distance of 1.0 neutral substitutions per site, an evolutionary time approximately proportional to 500 million years of vertebrate evolution (assuming a molecular clock of 0.002 neutral substitutions per site per million years). This process, which we term a burn-in, shuffles the sequences, genes and chromosomes of the genome. The resulting genome was termed the most recent common ancestor (MRCA) because it was used as the starting point for both the primate and mammalian simulations. It has been previously ascertained that distributions on the numbers and lengths of tracked annotation types in EVOLVER simulations stay stationary over time (Earl et al. 2011), so this burn-in process, from a simulation point of view, does not adversely affect the nature of the simulated genomes.

The primate simulation was described by the phylogenetic tree (in newick format; Fig. 1):

> ((simGorilla:0.008825,(simHuman:0.0067,simChimp:0.006667)sHuman-sChimp:0.00225)sG-sH-sC:0.00968,simOrang:0.018318);

The mammal simulation was described by the phylogenetic tree (in newick format; Fig. 1):

> ((simCow:0.18908,simDog:0.16303)sCow-sDog:0.032898,(simHuman:0.144018,(simMouse:0.084509,simRat:0.091589)sMouse-sRat:0.271974)sH-sM-sR:0.020593);

We used the EVOLVER produced repetitive element library from the simHuman genome as an input library for RepeatMasker. Following each simulation, the EVOLVER mobile element library from the simHuman leaf node genome was used as an input into the repetitive sequence finder RepeatMasker. RepeatMasker was then used to mask simple repeats and repeats from the provided library in the other non-human simulated genomes.

Complete sequence and annotations of the leaf genomes and the MRCA genome were provided to participants.

### Flies Dataset

The phylogeny was created by merging the phylogeny provided in the mod ENCODE comparative genomics white paper (http://www.genome.gov/Pages/Research/Sequencing/SeqProposals/modENCODE_ComparativeGenomics_WhitePaper.pdf accessed 15 October 2013) courtesy of Artyom Kopp (UC Davis) and the phylogeny used by UCSC for the 15-way insect alignment. The Kopp tree lacked droSim1 and droSec1 which were added by normalizing the branch lengths between the dm3 branches on the two trees. Extraneous species were trimmed using tree_doctor from PHAST. This tree was provided for progressive aligners that need a guide tree. This phylogeny corresponds to the newick tree (Fig. 1):

> ((droGri2:0.183954,droVir3:0.093575):0.000000,(droMoj3:0.110563,((((droBip:0.034265,droAna3:0.042476):0.121927,(droKik:0.097564,((droFic:0.109823,(((dm3:0.023047,(droSim1:0.015485,droSec1:0.015184):0.013850):0.016088,(dro Yak2:0.026909,droEre2:0.029818):0.008929):0.047596,(droEug:0.102473,(droBia:0.069103,droTak:0.060723):0.015855):0.005098):0.010453):0.008044,(droEle:0.062413,droRho:0.051516):0.015405):0.046129):0.018695):0.078585,(droPer1:0.007065,dp4:0.005900):0.185269):0.068212,droWil1:0.259408):0.097093):0.035250);

To create the fly sequence dataset we took 12 flies available from the UCSC golden path server on 14 December 2011 (droAna3, dreEre2, droGri2, droMoj3, dp4, droVir3, droWil1, dm3, droSim1, droPer1, droSec1, droYak2) and eight flies from NCBI on 25 January 2012 (droBia, droBip, droEle, droFic, droKik, droTak, droRho, droEug).

### MafTools

Participants submitted their predictions of alignments in MAF files. To process the submissions we wrote a suite of open-source tools called mafTools, available at https://github.com/dentearl/mafTools/ to perform the majority of transformations, manipulations and analyses. Scripts to perform the analyses described can be found in the analysis repository.

### MAF Comparisons

Exhaustively checking all pairs of aligned residues between alignments is computationally impractical, so instead we developed a method, termed *mafComparator*, based upon sampling pairs of aligned residues. Sampling is performed by reading each input MAF file twice, once to count the total number of pairs present in the file, such that given a user specified number of pairs to sample we can calculate the probability of picking a given pair at random. The MAF file is then read a second time.

During the second pass we iterate over every block in the MAF and then every column in the block. We calculate the number of pairs present in the block, call this *k*, and then make a draw from a binomial distribution with probability *s* / *m* (where s is the number of samples taken, here 10,000,000, and *m* is the total number of pairs present in the MAF) to see how many (if any) pairs to sample from that column. If *x* many pairs are to be sampled we then sample *x* times from a discrete uniform [0, *k* − 1] decrementing the range of the distribution with each sample, without replacement, and then map those integers to pairs using a bijective function. This allows us to efficiently sample pairs without iterating through each and every pair.

### Regional Alignments

To accommodate PSAR, which processes global MSAs in which the alignment is represented as a 2D matrix where the aligned sequences, interspersed with gaps, are the rows and the columns represent the equivalence classes of aligned bases, we constructed sub-alignments of sampled regions.

For each of the three test sets, regional intervals were selected by sampling five different starting values from a discrete uniform distribution [0, *g* - 1 - 500,000], where *g* is the total length of the reference genome and 500,000 is the length of the interval. Sampled values were then mapped back to individual chromosomes. All alignments containing any positions of the reference within these intervals were extracted from the submitted alignments. Though this model of sampling does not prevent overlapping regions, no overlapping regions were sampled. Likewise this model of sampling does not prevent regions that cross between chromosomes, but no such bridged regions were sampled.

To convert each subregion into a 2D alignment matrix we designed a pipeline that used the chosen reference genome interval to create a “reference alignment” as follows:

1. For each submission, for each region, we used mafExtractor to pull out maf blocks that contained any positions in the reference that were within the target region. These blocks were then trimmed to contain only positions that aligned to the reference within the region of interest.
2. We used our mafTransitiveClosure tool to compute the transitive closure of the resulting alignment (see discussion), to make each alignment a consistent multiple sequence alignment. This was particularly important for the GenomeMatch alignments, which were all pairwise. Unlike for the whole genome alignments, taking the transitive closure of the aligned pairs just within the subregion did generally not perturb the performance substantially (data not shown).
3. Next, a row deduplication step was performed using mafDuplicateFilter, so that each species contributed at most one row to the 2D alignment. In short, for every block in the alignment a consensus sequence is created and then for species with multiple instances present in the block a similarity score is calculated in relation to the consensus. Only the sequence closest to the consensus in terms of substitution distance is kept. In the event of a tie the sequence closest to the start of the block is used. All the other instances are discarded.
4. The previous step could result in the removal of the reference sequence that tied the block into the region of interest. As a result, a second step of reference oriented region based extraction is performed to eliminate all blocks that more strongly align to an area outside of the region of interest (i.e. a part of the reference outside the region of interest had a higher similarity score in the row deduplication step).
5. The blocks are sorted based upon their left-right order along the positive strand of the region of interest in the reference. Then the rows of all blocks are reordered to be standardized (into a consistent species order, i.e. alphabetically) and finally all of the reference sequence instances are forced to be in relation to the positive strand. Rearrangements in the non-reference sequences were ignored by concatenating the fragments of the non-reference sequence together according to their ordering along the reference sequence, as is standard practice in constructing “reference” alignments.

In calculating the regional precision and recall values for the simulated subregions we ignored all pairs in both the true alignment and the predicted subregion alignment that did not contain a residue from the sampled reference species (sub)sequence, i.e. we computed an analogous regional version of these scores.

## Acknowledgements

We would like to thank the Howard Hughes Medical Institute, Dr. and Mrs. Gordon Ringold, NIH grant 2U41 HG002371-13 and NHGRI/NIH grant 5U01HG004695 for providing funding. We would like to thank the Genome 10K organizers for providing a venue to discuss an early version of these findings.

